# Differential expression of P2RY12 on the protein level between adenocarcinomas and squamous cell carcinomas reveals potential diagnostic value and a possible guidance for anti-P2RY12 therapies in NSCLC

**DOI:** 10.64898/2026.06.13.732072

**Authors:** Christiane Kuempers, Hauke Roettger, Tobias Jagomast, Lukas Emken, Carsten Heidel, Finn-Ole Paulsen, Tristan Tuecking, Jutta Kirfel, Daniel Droemann, Sabine Bohnet, Michael Schweigert, Martin Reck, Till Olchers, Soenke von Weihe, Volker Meidl, Doerte Nitschkowski, Torsten Goldmann

## Abstract

**Context:** P2Y12 receptor (P2RY12), mainly expressed on platelets, is known for its central role in hemostasis. P2RY12 activation is also involved in cancer development through platelet adhesion to cancer cells, supporting immune evasion, promoting tumor angiogenesis and metastasis, among others. P2RY12 is known as an actionable target, and P2RY12 antagonists are in clinical use for cardiovascular diseases. There is very little data available regarding the protein expression of P2RY12 in lung carcinomas.

**Design:** We performed immunohistochemical staining for P2RY12 in a cohort of NSCLC samples (319 primary adenocarcinomas (LUAD) and 157 primary squamous cell carcinomas (LUSC)). Results were evaluated using a dual approach comprising microscopic assessment and digital image analysis (QuPath). Results were correlated with clinical-pathological data. Results: We found significantly higher P2RY12 protein expression in LUSC than in LUAD (p<0.001) via eyeballing (absent/low expression in 21.7% (34/157) and moderate/high expression in 78.3% (123/157) of LUSC cases versus absent/low expression in 98.4% (314/319) and moderate/high expression in 1.6% (5/319) of LUAD cases). Digital analysis yielded similar results. Kaplan-Meier analysis showed numerically higher 5-year overall survival probabilities in patients with high P2RY12 expression, although the differences were not statistically significant. However, the discriminatory performance of P2RY12 for classifying carcinomas as LUAD or LUSC was excellent (AUC of 0.916 in ROC analysis).

**Conclusion:** Although P2RY12 expression was not significantly associated with overall survival and was not identified as an independent factor in multivariable analysis, the numerically higher survival probabilities observed in patients with high P2RY12 expression warrant further investigation of its potential prognostic and therapeutic relevance. Importantly, the pronounced differences in P2RY12 expression between LUSC and LUAD suggest promising discriminatory potential, which requires validation in independent cohorts.

## Introduction

The two principal forms of lung cancer are non-small cell lung cancer (NSCLC), accounting for approximately 85% of all cases, and small cell lung cancer (SCLC), comprising the remaining 15% (Alduais et al., 2023). Among NSCLC subtypes, lung adenocarcinoma (LUAD) (50%) and lung squamous cell carcinoma (LUSC) (20-30%) are the most prevalent histological entities (Hendriks et al., 2024). A deeper understanding of molecular drivers and tumor biology is essential to close existing gaps in treatment and prevention (Y. Li et al., 2023). Although advances in systematic screening, the introduction of novel surgical and radiotherapeutic approaches, and improved biological understanding of NSCLC have contributed to better outcomes (Howlader et al., 2020), lung cancer remains the leading cause of cancer-related mortality worldwide (Bray et al., 2024).

Important drivers of mortality in cancer are cancer-associated thrombosis and the development of distant metastases, with platelets potentially playing a pivotal role at their intersection (Falanga & Marchetti, 2023). Cancer cells can induce platelet activation via FcγRIIa, a low-affinity IgG receptor expressed on the platelet surface, triggering the release of bioactive molecules such as lipids and growth factors, which facilitate platelet–tumor cell adhesion (Bellefeuille et al., 2019). Through surface receptors like integrins, platelets form complexes with tumor cells, shielding them from immune clearance by natural killer (NK) cells and tumor necrosis factor alpha (TNF-α) (Gresele et al., 2018).

Additionally, platelets release vascular endothelial growth factor (VEGF) and other pro-angiogenic factors, supporting tumor vascularization (Radziwon-Balicka et al., 2012). Furthermore, platelet-derived microparticles may transfer bioactive cargo, such as messenger ribonucleic acid (mRNA), microRNA, non-coding RNAs, proteins, receptors, and mitochondria, to other cells, including stromal, immune, epithelial, and cancer cells, modulating their phenotype and promoting tumor progression and metastasis (Dovizio et al., 2018).

Inhibition of platelet activation may therefore represent a therapeutic approach in cancer, with the potential to reduce both metastatic spread and thrombotic risk (Wright et al., 2020). The P2Y12 receptor (P2RY12) represents one of the principal regulators of platelet activation and is crucial for platelet function, hemostasis, thrombosis (X. Li et al., 2023), and the pathogenesis of inflammation-related diseases (Parker & Storey, 2024). At present, there are eight recognized human P2Y receptors (Burnstock, 2018). This study focuses on the purine receptor P2RY12 subtype. P2RY12 has emerged as a central therapeutic target in cardiovascular medicine, with P2RY12 inhibitors, such as clopidogrel, prasugrel, and ticagrelor, now firmly established as components of standard care in the prevention and treatment of atherothrombotic events (Patti et al., 2020).

P2RY12 is a G-protein coupled receptor with seven transmembrane domains and is activated by extracellular nucleotides, which function as signaling molecules in various physiological and pathological processes (Cattaneo, 2015).

P2RY12 expression is not restricted to platelets but has been identified in multiple physiological cells and tissues, including distinct regions of the brain, where it serves as a specific marker for microglial cells (Mildner et al., 2017), as well as in subsets of immune cells (Entsie et al., 2023), vascular smooth muscle cells, and endothelial cells (Wihlborg et al., 2004). Beyond its physiological distribution, P2RY12 expression has been documented in pancreatic cancer cell lines (Elaskalani et al., 2020), breast cancer cell lines (Sarangi et al., 2013), hepatocellular carcinoma mouse models (Ma et al., 2022), and glioblastoma cell lines (Vargas et al., 2022). Regarding lung carcinoma, experimental studies in Lewis Lung Carcinoma (LLC) spontaneous metastatic mouse models demonstrated that genetic deficiency of P2RY12 led to attenuated tumor growth and a significant reduction in metastatic dissemination. Their results further indicated that P2RY12 deficiency diminished the ability of LLC cells to induce platelet shape change and release of active TGFβ1, resulting in a diminished, platelet-induced epithelial-mesenchymal transition (EMT)-like transformation of the LLC cells as a probable prerequisite of LLC cell metastasis (Y. Wang et al., 2013).

Consistent with these preclinical observations, an *in vitro* investigation demonstrated that ticagrelor, a clinically used antagonist of the platelet P2RY12, inhibited proliferation and colony formation as well as migration and invasion in LUAD cells. The authors reported similar findings in *in vivo* analyses with lung adenocarcinoma cell-bearing mouse models demonstrating inhibited growth of lung cancer xenografts following Ticagrelor administration (Song et al., 2025). To our knowledge, the sole available immunohistochemical analysis of resected tumor specimens from 37 patients with lung adenocarcinoma revealed that low P2RY12 expression was significantly associated with an unfavorable prognosis (Yu et al., 2021).

Despite promising evidence from previous studies, data on P2RY12 protein expression in human NSCLC specimens remain scarce. This study therefore aimed to analyze P2RY12 protein expression in a large NSCLC study population and to assess its correlation with survival data and other clinico-pathological parameters.

## Material and Methods

### Study Design and Patient Population

The study was approved by the ethics committee of the University of Luebeck (project code AZ 16-277 and 16-278) and was conducted in accordance with the principles of the Declaration of Helsinki. A total of 535 patients with histologically confirmed NSCLC underwent surgical resection. Of these, 363 patients (67.9%) were diagnosed with LUAD, 168 (31.4%) with LUSC, and 4 (0.7%) with NSCLC not otherwise specified (NOS). After excluding these 4 NOS cases and an additional 17 samples that were not evaluable due to technical artifacts or tissue loss during TMA construction, a total of 514 tumor samples remained evaluable for further analysis.

Of these, 478 were primary tumors (319 LUAD, 157 LUSC, 2 without histological subtype information), out of which 38 had matched metastatic samples, while 36 patients contributed metastatic samples only (3 lymph node and 33 distant metastases) (Figure 1). Because histological subtype information was missing for two of these samples, 476 primary tumors (319 LUAD, 157 LUSC) were available for all analyses stratified by histological subtype (comparison of histological subtypes, tissue comparison and ROC analysis).

**Figure 1.**
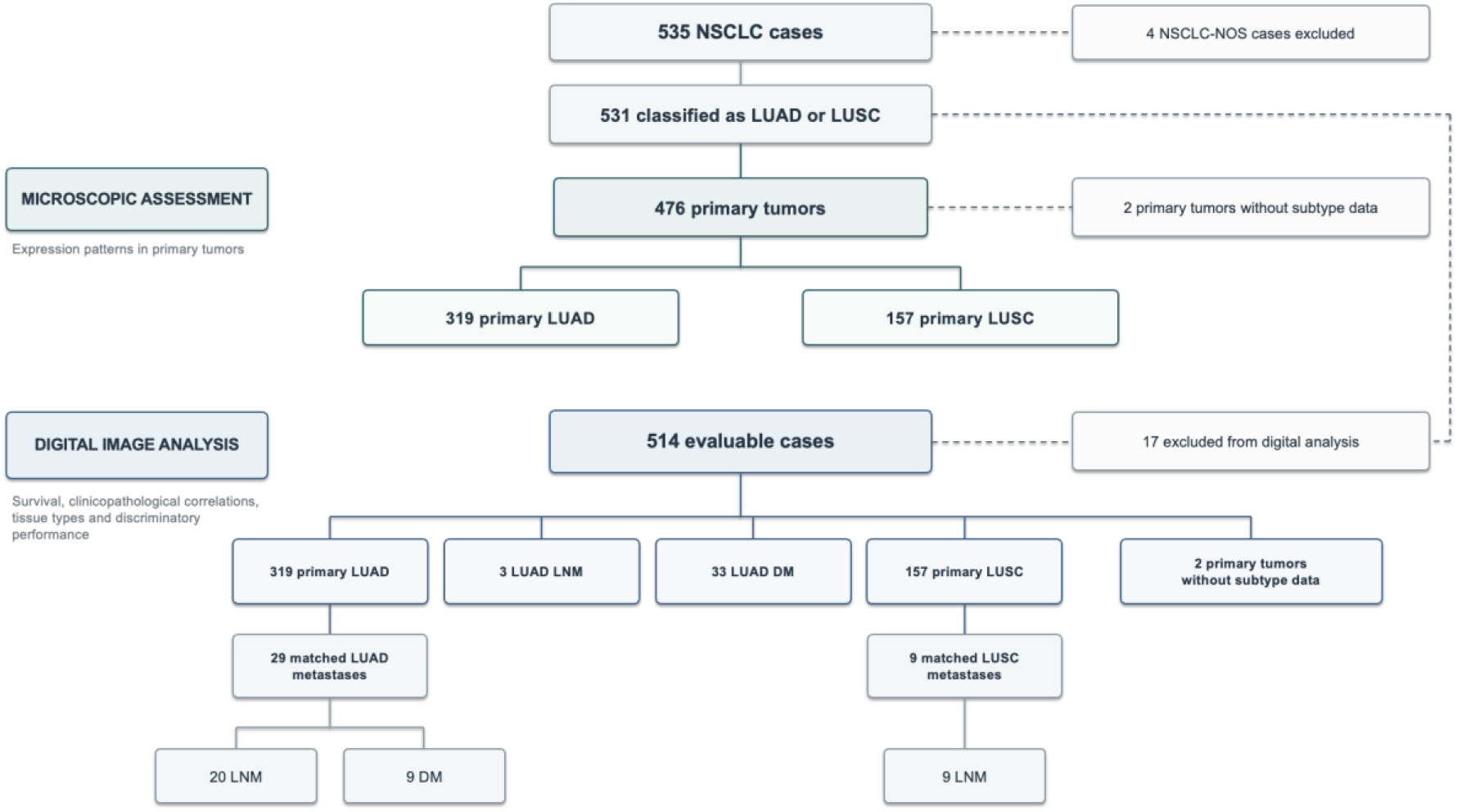
Flowchart of the cohort illustrating each analytical subgroup. The flowchart summarizes cohort selection and composition. The upper green section represents the cohort of primary tumors for the microscopic assessment, and the lower blue section shows the cohort for subsequent digital image analysis including metastases additionally. [LNM: lymph node metastases; DM: distant metastases]

At the time of the last follow-up, 338 patients (63.2%) were alive, 120 (22.4%) were deceased, and for 77 patients (14.4%) follow-up data were unavailable.

Clinicopathological data were analyzed for 514 evaluable cases. Pathological T stage and histological grading were available for 511 and 495 cases, respectively. In the entire cohort of 535 patients, missing data were observed for age (n=5), pT stage (n=24), pN status (n=66), M stage (n=401), UICC stage (n=31), tumor grade (n=40), and recurrence status (n=61). Furthermore, valid follow-up time was unavailable for 120 patients.

For survival analyses, 366 of 478 biomarker-evaluable patients (76.6%) had sufficient follow-up information. The number of patients included in each analysis is reported separately in the corresponding results and supplementary table.

The clinico-pathological characteristics of the study population are depicted in Table 1. Figure 1 shows the cohort with each analytical subgroup.

**Table 1.**
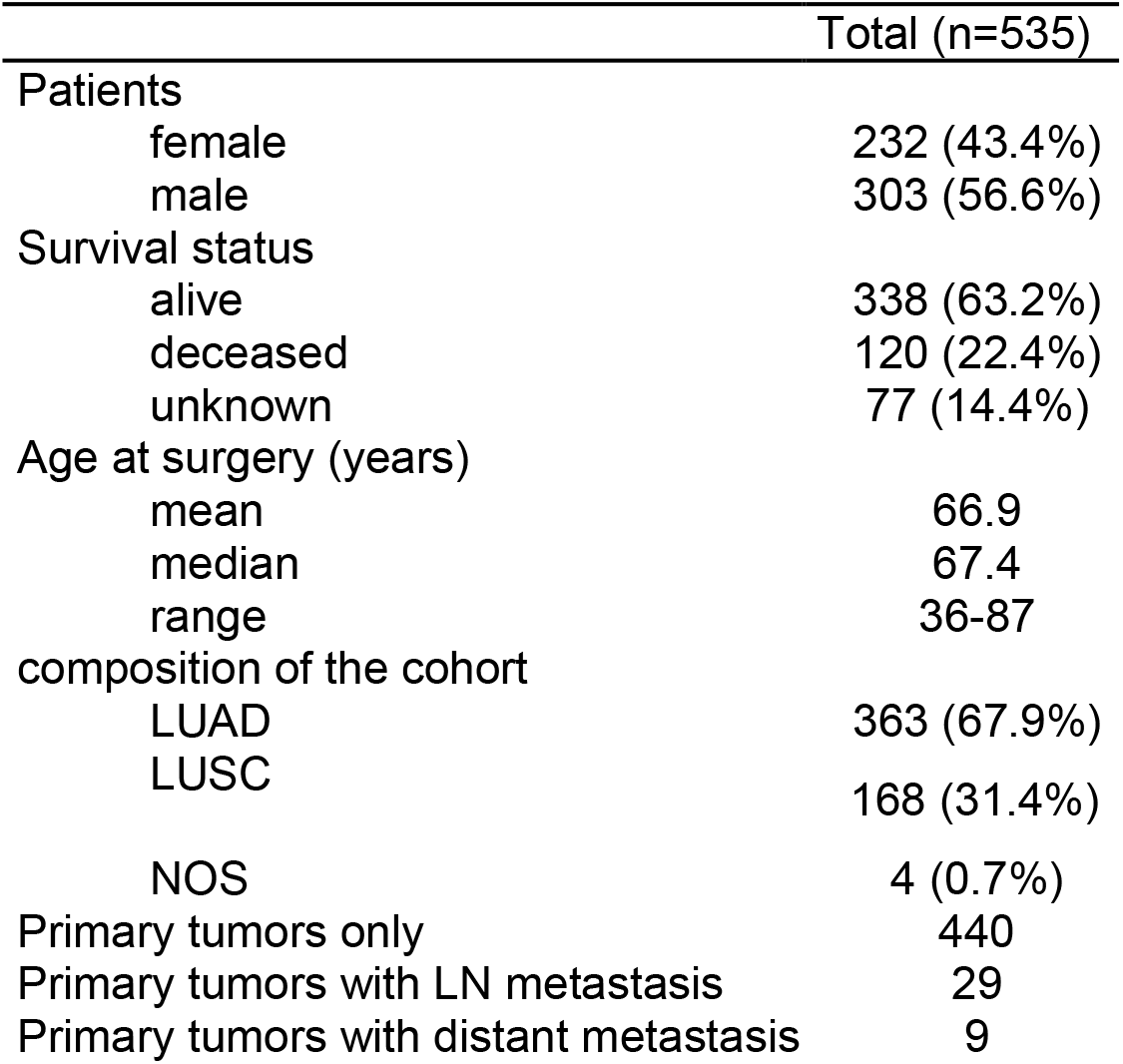
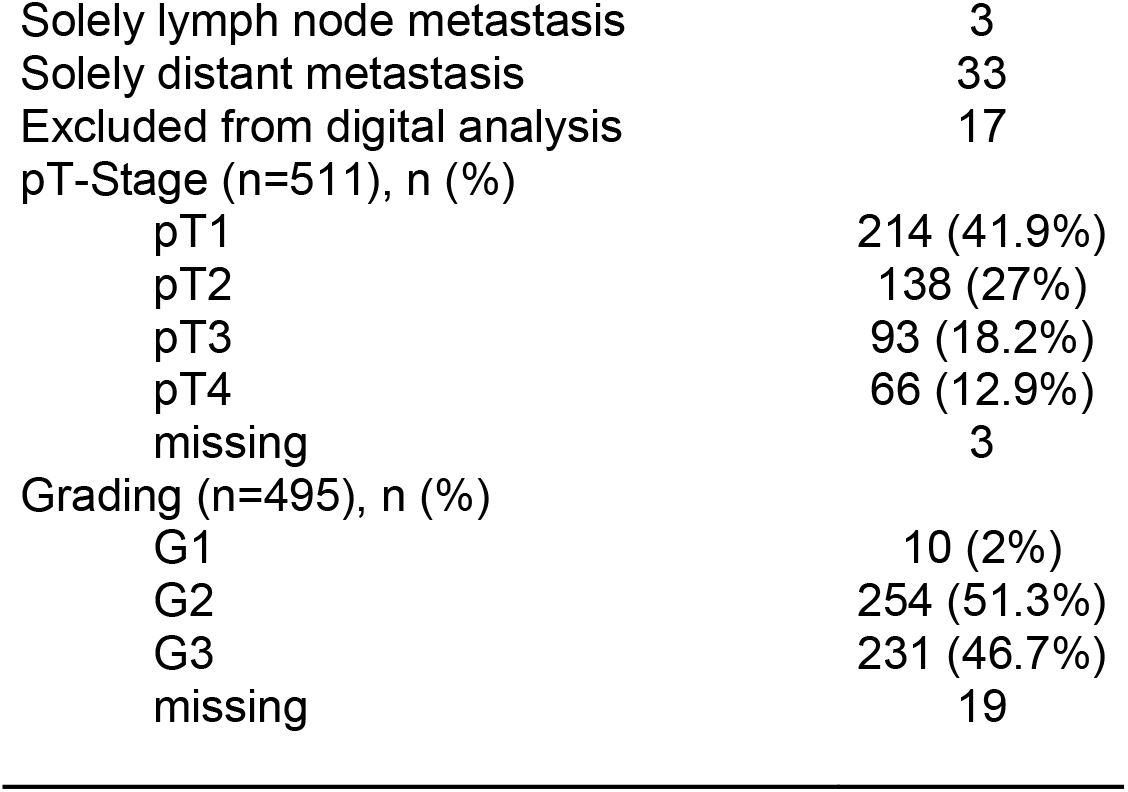
Clinico-pathological characteristics of the cohort. Tumor grading was performed according to the 2015 World Health Organization (WHO) Classification of Lung Tumors (Travis et al., 2015). Pathological T-staging was assigned according to the 8th edition of the Union for International Cancer Control (UICC) TNM classification (Lim et al., 2018).

|  | Total (n=535) |
|---|---|
| Patients |  |
| female | 232 (43.4%) |
| male | 303 (56.6%) |
| Survival status |  |
| alive | 338 (63.2%) |
| deceased | 120 (22.4%) |
| unknown | 77 (14.4%) |
| Age at surgery (years) |  |
| mean | 66.9 |
| median | 67.4 |
| range | 36-87 |
| composition of the cohort |  |
| LUAD | 363 (67.9%) |
| LUSC | 168 (31.4%) |
| NOS | 4 (0.7%) |
| Primary tumors only | 440 |
| Primary tumors with LN metastasis | 29 |
| Primary tumors with distant metastasis | 9 |
| Solely lymph node metastasis | 3 |
| Solely distant metastasis | 33 |
| Excluded from digital analysis | 17 |
| pT-Stage (n=511), n (%) |  |
| pT1 | 214 (41.9%) |
| pT2 | 138 (27%) |
| pT3 | 93 (18.2%) |
| pT4 | 66 (12.9%) |
| missing | 3 |
| Grading (n=495), n (%) |  |
| G1 | 10 (2%) |
| G2 | 254 (51.3%) |
| G3 | 231 (46.7%) |
| missing | 19 |

### Immunohistochemistry (IHC)

Tissue microarrays (TMAs) were constructed from formalin-fixed, paraffin-embedded (FFPE) tumor blocks from resection specimens. All cases were therapy-naïve. For each case, at least three 1.2 mm diameter cores were obtained from representative tumor regions and transferred into recipient paraffin blocks using a manual tissue arrayer. To reduce the likelihood of systematic staining artifacts, at least three representative tumor cores were sampled per case, and tumors were considered evaluable if at least one core yielded sufficient tissue. In total, 17 TMAs were constructed (12 LUAD, 5 LUSC), allowing cross-validation across independent TMAs.

To perform the immunohistochemical staining, formalin-fixed, paraffin-embedded (FFPE) lung tumor sections were deparaffinized, rehydrated, and subjected to heat-induced epitope retrieval in citrate buffer (pH 6.0; 30 minutes at 95 °C, steam cooker). Endogenous peroxidase activity was inhibited with 3% hydrogen peroxide for 10 minutes at room temperature. Sections were incubated for 60 minutes at room temperature on a shaker with a polyclonal rabbit anti-P2RY12 antibody (clone: #APR-012, Alomone labs; dilution 1:1000). Detection was performed using a horseradish peroxidase (HRP)-polymer (anti-mouse/rabbit; 20 minutes, room temperature), followed by development with 3-amino-9-ethylcarbazole (AEC) chromogen for 10 minutes. Sections were counterstained with Mayer’s hematoxylin (1:3 dilution) for 5 seconds, blued in 0.05% ammonia water for 20 seconds, and finally dehydrated through a graded ethanol series, cleared in xylene, and mounted using Pertex. A negative control without the primary antibody was performed to exclude nonspecific background staining resulting from the detection system or other components of the staining procedure (Supplement Figure 1).

### Microscopic Evaluation

Following immunohistochemical staining of the TMAs, all patient samples were independently evaluated by two investigators using a light microscope (Olympus BH-2). Staining intensity was categorized into four levels: absent, weak, moderate, and strong. In cases of substantial disagreement, the respective samples were jointly reassessed until a consensus score was reached. Interobserver agreement was not formally assessed. For dichotomous analyses, specimens with absent or weak staining were classified as low, whereas those with moderate or strong staining were classified as high. Exemplary pictures are shown in Figure 3.

### Digitalization

Slides were additionally digitized with a Ventana DP 200 scanner (Roche), and TIF files were generated. Image analysis was performed with QuPath (Quantitative Pathology, version 0.5.1, https://qupath.github.io), an open-source bioimage analysis platform enabling automated quantification of staining intensities within user-defined regions of interest (ROIs). Scanned TMA images were imported into QuPath, and individual tissue cores were assigned to the corresponding patients using TMA grids. Representative areas were manually annotated as tumor, stroma, immune cells, or anthracosis and used to train the classifier. Cell detection was performed using the optical-density-sum method at a pixel size of 0.5 µm. Cell nuclei were detected using a background radius of 5 µm, a median filter radius of 1 µm, a sigma of 1 µm, a nuclear area of 10 to 500 µm², and an intensity threshold of 0.08. Cell boundaries were defined by a 6-µm expansion beyond the nucleus. Smoothed features with a radius of 25 µm were additionally incorporated to improve classification accuracy.

Staining intensity was quantified by optical density (OD). Given the cytoplasmic localization of P2RY12 staining, the mean cytoplasmic OD was automatically calculated for each core. Thresholds were established by conventional microscopic evaluation and categorized as negative (OD, 0–0.13), weak (0.13–0.25), moderate (0.25–0.47), or strong (>0.47). Cells were subsequently assigned to tissue classes using the integrated QuPath classifier with a random-forest algorithm trained on the manually annotated cells from the entire project, with balanced representation of tumor, stromal, and immune cells. The classifier combined multiple decision trees trained on random subsets of observations and features to distinguish cell classes based on morphological and staining characteristics. The resulting classifier was applied to all samples for automated identification of regions of interest and classification of detected cells (Figure 2). For each patient, the mean staining intensity of cells classified as tumor was calculated using a custom Groovy script and exported for subsequent statistical analysis.

**Figure 2.**
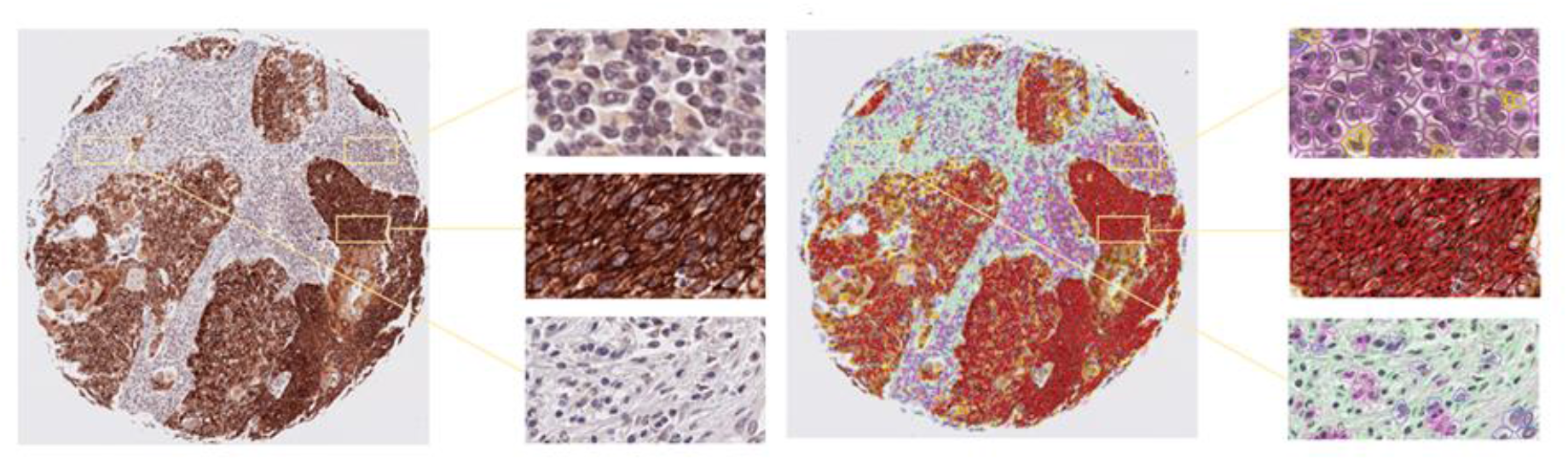
TMA core without (left) and with (right) segmentation of immune cells (top), tumor (middle), and stroma (bottom)

TMA cores were excluded from the microscopic and digital analysis if they contained no or insufficient tumor tissue, showed necrosis, or had detached from the slide. Cores were also excluded from digital analysis when tissue folds or air bubbles beneath the coverslip caused scanning and focus artefacts that precluded reliable tissue classification on QuPath. Non-evaluable samples were excluded from statistical analyses irrespective of P2RY12 expression, histologic subtype, or clinical outcome, since reliable P2RY12 expression values could not be obtained.

A case was included if at least one of the three cores was evaluable. Data acquisition was performed on a Windows 11 Pro workstation with a 27-inch monitor (resolution 2560 × 1440 pixels).

### Statistical analyses

All statistical analyses were performed with R (version 4.4.2; R Foundation for Statistical Computing, Vienna, Austria) using the packages survival (3.7-0), survminer (0.5.0), arsenal (3.6.3), pROC (1.18.5), mice (3.17.0) and car. All tests were two-sided, and p-values <0.05 were considered statistically significant. Subgroup and clinico-pathological analyses were exploratory; unless stated otherwise, no adjustment for multiple comparisons was performed. Each analysis included all patients with the data required for that analysis (available-case analysis), and the number of patients analyzed is reported for each analysis. Because P2RY12 staining intensities (OD values) showed a right-skewed, non-normal distribution with a second peak formed by LUSC, non-parametric tests were used for all comparisons of staining intensity.

Agreement between the ordinal visual score (absent, weak, moderate, strong) and the continuous OD values was assessed with Spearman rank correlation, which requires neither a linear relationship nor normally distributed data; the 95% confidence interval was obtained by bootstrapping (2,000 replicates). Differences in OD values across the four visual categories were tested with the Kruskal–Wallis test.

The dichotomized visual score (absent/weak vs. moderate/strong) was compared between LUAD and LUSC with Pearson’s chi-square test, which was appropriate because all expected cell counts were large (minimum 42).

OD values of primary LUSC, primary LUAD, lymph node metastases and distant metastases were compared with the Kruskal–Wallis test followed by pairwise Wilcoxon rank-sum tests; pairwise p-values are additionally reported after Holm adjustment. Because primary tumors and their matched metastases are dependent observations, paired samples were compared with the Wilcoxon signed-rank test; for patients with both lymph node and distant metastases, the lymph node metastasis was used. The association between OD values and dichotomous clinico-pathological parameters (sex, pT, pN, M and UICC stage, grading, recurrence) was assessed with the Wilcoxon rank-sum test, and the association with age with Pearson correlation (Spearman correlation as sensitivity analysis).

For dichotomous analyses, P2RY12 expression was classified as high or low using the median OD value of all primary tumors (OD 0.141) as a pre-specified cut-off that was not optimized on outcome data. Clinico-pathological characteristics of the high- and low-expression groups were compared with Fisher’s exact test for categorical variables, because some cells (e.g., M1 or UICC stage IV) contained very small numbers, and with the Wilcoxon rank-sum test for continuous variables.

Overall survival was calculated from the date of diagnosis to death or last follow-up and was restricted to 5 years (administrative censoring at 260 weeks); death from any cause was the event, and living patients were censored at their last follow-up. Patients with unknown vital status or missing follow-up dates could not be included. Survival curves were estimated with the Kaplan–Meier method and compared with the log-rank test, for the entire cohort and separately for LUAD and LUSC; unadjusted hazard ratios (HR) were estimated with univariable Cox regression. Median follow-up was estimated with the reverse Kaplan–Meier method. Sensitivity analyses used histology-specific median cut-offs and the visual score (moderate/strong vs. absent/weak) instead of the cohort median of the OD values. To assess whether missing follow-up was related to the biomarker, patients with and without follow-up information were compared (Fisher’s exact test, Kruskal–Wallis test).

The independent prognostic value of P2RY12 expression was assessed with multivariable Cox proportional hazards regression. Covariates were pre-specified on clinical grounds and not selected by p-values: age (continuous, per year), sex, histology, pN status, tumor grade and pT stage. M stage was not included because it was missing in 79% of the biomarker-evaluable patients, and UICC stage was not included because it is derived from T, N and M and is therefore collinear with these variables. The proportional hazards assumption was assessed with scaled Schoenfeld residuals. Because the assumption was violated for pT stage when pT was entered as a covariate, the primary model was stratified by pT stage (T1–2 vs. T3–4), which allows separate baseline hazards for both strata; consequently, no HR is estimated for pT. Linearity of continuous covariates was checked with martingale residuals, collinearity with variance inflation factors (VIF), and the number of events per estimated parameter is reported. Model discrimination was assessed with Harrell’s concordance index. The primary model was a complete-case analysis. Sensitivity analyses comprised (i) multiple imputation of missing covariates by chained equations (20 imputed data sets; event indicator and Nelson–Aalen cumulative hazard included as predictors; estimates pooled with Rubin’s rules), (ii) P2RY12 expression as a continuous variable (HR per 0.1 OD unit), (iii) a model additionally stratified by histology, because P2RY12 expression and histology are correlated, and (iv) separate models for LUAD and LUSC, which were interpreted with caution because of the lower number of events.

The ability of the continuous OD value to discriminate LUSC from LUAD was assessed with receiver operating characteristic (ROC) analysis; the area under the curve (AUC) is reported with 95% confidence intervals calculated with the DeLong method and with a stratified bootstrap (1,000 replicates). The optimal cut-off was determined with the Youden index, and sensitivity, specificity, positive and negative predictive values and accuracy are reported with exact (Clopper–Pearson) 95% confidence intervals. Because the cut-off was derived from the same data, its optimism was estimated by internal validation: a bootstrap procedure (1,000 replicates) in which the cut-off selection was repeated in each replicate and evaluated in the original data, and repeated stratified five-fold cross-validation (50 repeats) in which the cut-off was determined in the training folds only. Sensitivity and specificity of the visual score (moderate/strong vs. absent/weak) were calculated for comparison.

## Results

### P2RY12 expression pattern

P2RY12 demonstrated a cytoplasmic staining pattern, with homogeneous expression across tissue cores from the same tumor, indicating negligible intratumoral heterogeneity. Across tumor specimens, expression intensity ranged from absent to strong immunoreactivity in LUSC, whereas LUAD showed predominantly absent or weak staining (moderate in 4 and strong in 1 of 319 cases (0.3 and 1.3%, respectively)). Representative images of the various histological expression levels, including the corresponding OD values are shown in Figure 3. In addition, P2RY12 expression was observed in basal respiratory epithelium (n=14), with a representative example presented in Figure 3i).

**Figure 3.**
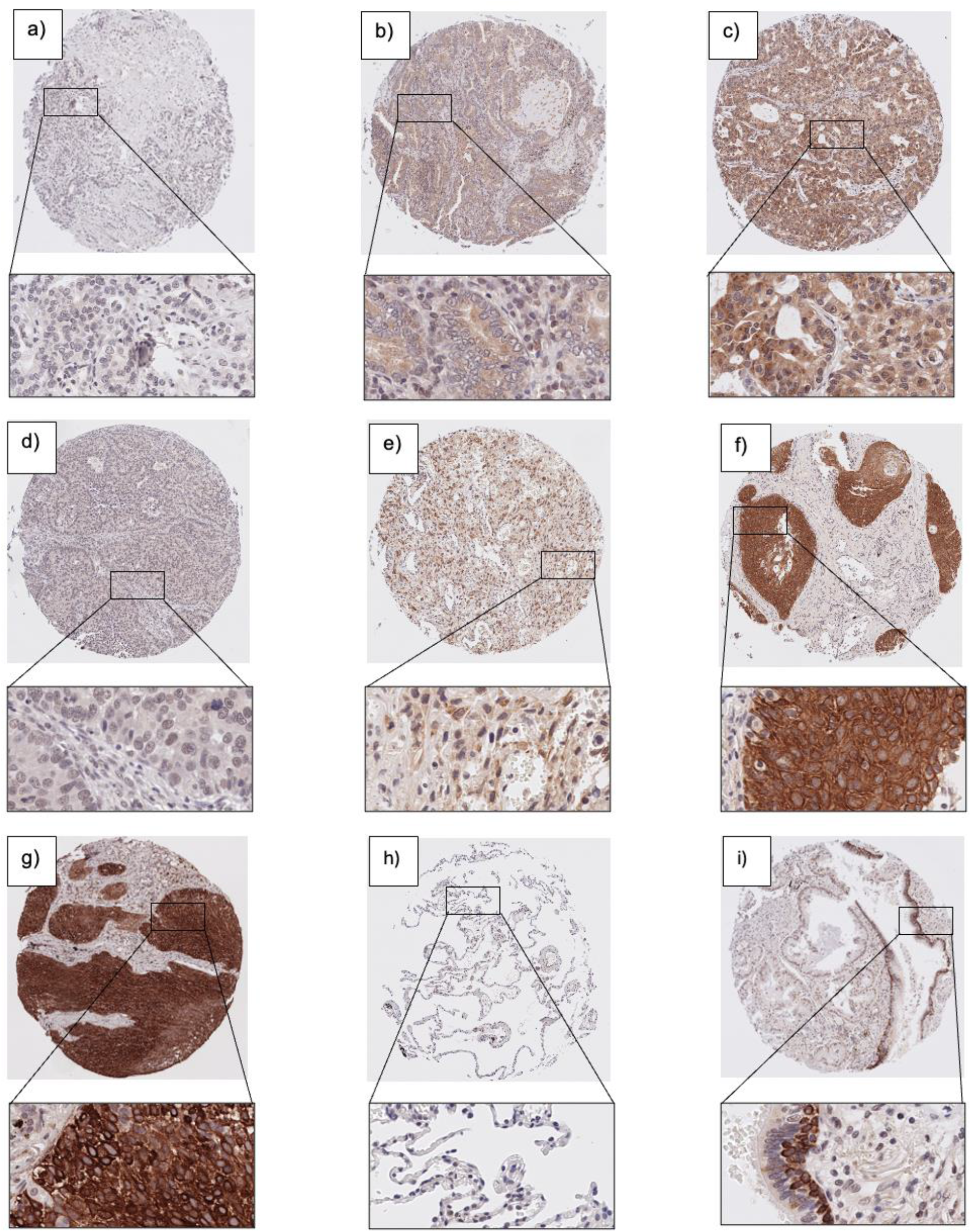
Exemplary pictures of P2RY12 expression pattern in NSCLC. ***a)-c)*** LUAD with a) absent (OD 0.08), b) low (OD 0.15), c) moderate (OD 0.26) expression of P2RY12. A high expression pattern was found in only one LUAD case. **d)-g)** LUSC with d) absent (OD 0.11), e) low (OD 0.16), f) moderate (OD 0.39), g) high (OD 0.64) expression of P2RY12. ***h)*** Normal tissue with negligible P2RY12 expression in pneumocytes. ***i)*** Normal tissue with P2RY12 expression in basal bronchial epithelium Objective Magnification: x 100 and x 400, respectively.

We first analyzed the distribution of eyeballing staining intensity of P2RY12 between LUAD and LUSC based on visual inspection. Among the 476 evaluable primary tumor samples with known histological subtype, marked differences in P2RY12 expression were observed between LUAD and LUSC. In LUAD, staining intensity was absent or low in 314 of 319 cases (98.4%), and only 5 cases (1.6%) showed moderate or high expression. In contrast, 123 of 157 LUSC cases (78.3%) demonstrated moderate or high expression, whereas absent or low expression was observed in 34 cases (21.7%). Overall, moderate or high expression was detected in 128 of 476 tumors (26.9%), the vast majority of which were LUSC (96.1%).

The distribution of expression levels differed significantly between histological subtypes of primary tumors (p<0.001, chi-square test) (Table 2). Moderate or high expression was almost exclusively confined to LUSC, whereas LUAD was characterized by absent or weak expression in nearly all cases.

**Table 2.** Contingency table of P2RY12 eyeballing staining intensity by histological subtype (n = 476 primary tumors)

|  |  | Eyeballing staining intensity |  |  |
|---|---|---|---|---|
|  |  | absent and low | moderate and high | Total |
| Histology | LUAD | 314 | 5 | 319 (67.0%) |
|  | LUSC | 34 | 123 | 157 (33.0%) |
|  | Total | 348 (73.1%) | 128 (26.9%) | 476 |

### Concordance between Computer-Assisted and Observer-Based Evaluation of P2RY12 Expression

At the outset, we compared the two evaluation methods statistically. To assess the validity of the computer-assisted results, we correlated visual scoring with quantitative measurements. A strong positive correlation was observed between visual scoring of P2RY12 staining intensity and quantitative optical density values obtained with QuPath (Spearman’s ρ = 0.78, 95% CI 0.74–0.82; p<0.001; n=478). Increasing visual categories (negative, low, medium, high) were associated with progressively higher median optical density values (0.11, 0.14, 0.30, and 0.44, respectively; Kruskal–Wallis test, p<0.001), indicating a high degree of concordance between subjective and objective evaluation methods (Figure 4).

**Figure 4.**
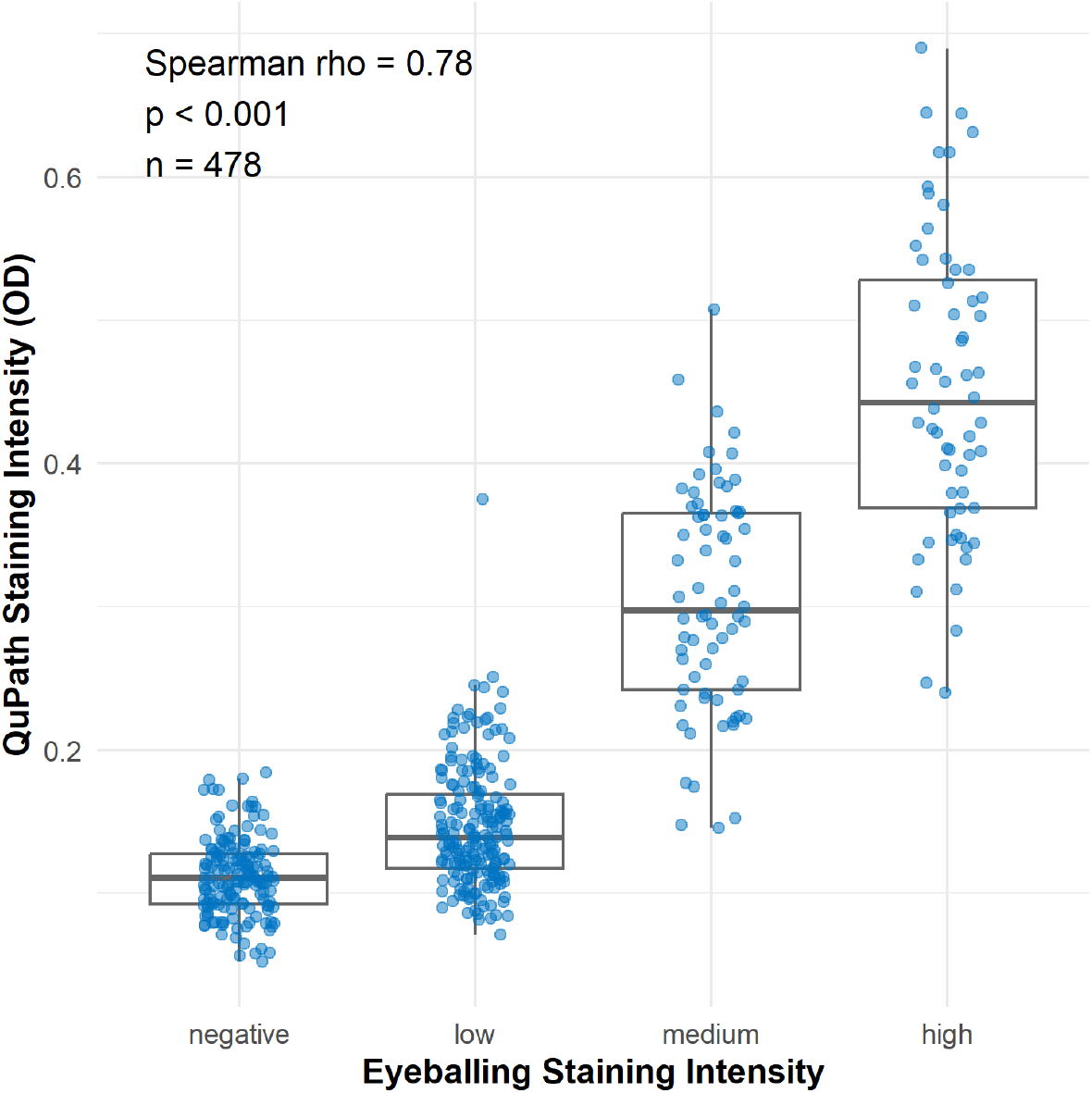
Scatter plot showing correlation between visual and computer-assisted assessment of P2RY12 staining.

Since the evaluation of staining patterns yielded equivalent results with both approaches, we used quantitative QuPath measurements for all subsequent analyses.

### Correlation of P2RY12 staining intensity across tissue origins

To compare staining intensity values across tissue origins, the Kruskal–Wallis test (p<0.001) followed by pairwise Wilcoxon rank-sum tests was applied (Figure 5A). Staining intensity was significantly higher in LUSC (n=157; median OD 0.34) compared with all other tissue groups: LUAD (n=319; median OD 0.12), lymph node metastases (LNM) (n=34 from 25 LUAD and 9 LUSC; median OD 0.12), and distant metastases (DM) (n=42, all from LUAD; median OD 0.11; all p<0.001, also after Holm adjustment). In contrast, no significant differences in staining intensity were detected among LUAD, LNM, and DM samples (p=0.35 for LUAD vs. LNM; p=0.31 for LUAD vs. DM; p=0.93 for LNM vs. DM; Holm-adjusted p=0.93 for each of these comparisons). Thus, strong expression of P2RY12 was largely confined to LUSC, whereas LUAD, LNM, and DM displayed uniformly lower staining levels.

**Figure 5.**
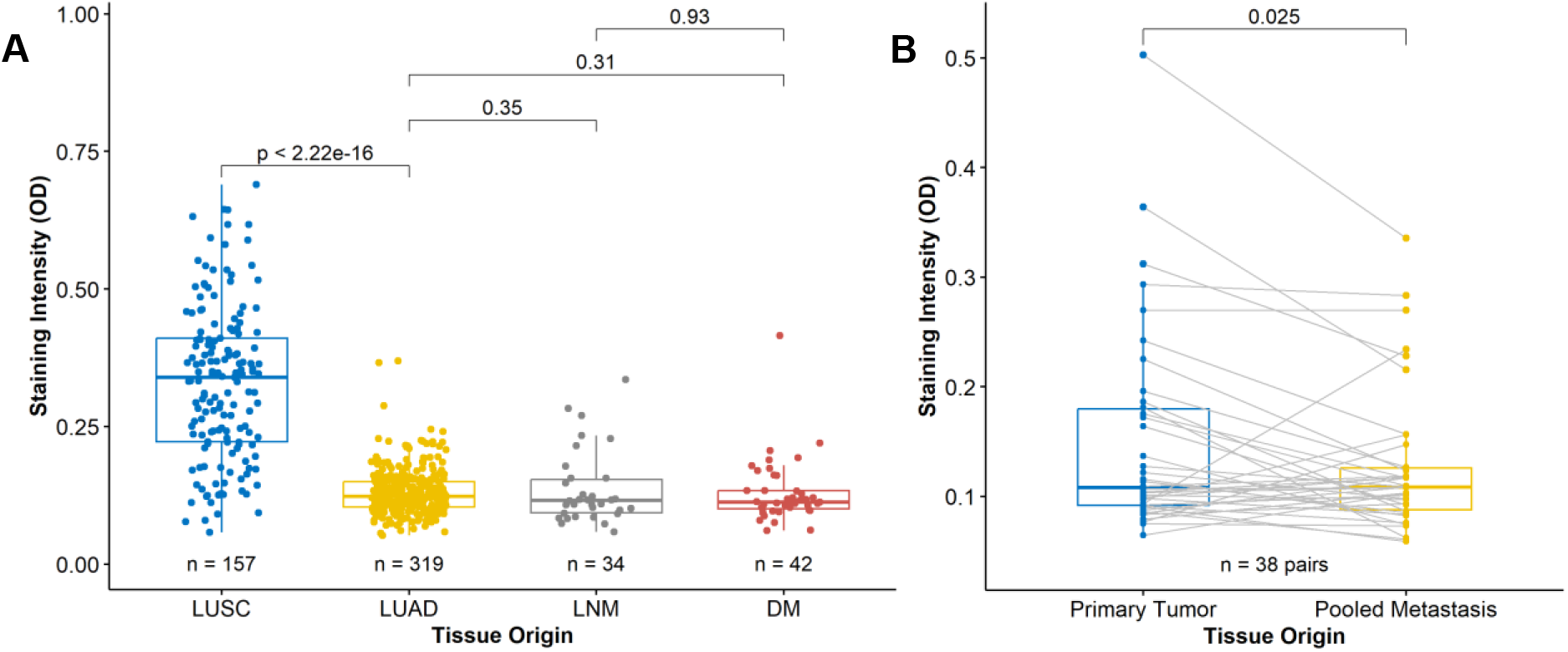
(A) Box plots show the distribution of P2RY12 staining intensity across primary LUAD and LUSC, as well as lymph node metastases [LNM] and distant metastases [DM]; (B) Box-and-whisker plots display the distribution of OD values in 38 paired samples of primary tumors (blue) and pooled metastases (yellow)

In the analysis of 38 paired samples (LNM and DM, from 29 LUAD and 9 LUSC), staining intensity of P2RY12 was modestly but significantly (median paired difference −0.015 OD; Wilcoxon signed-rank test, p=0.025) lower in metastases compared with the corresponding primary tumors (Figure 5B).

P2RY12 expression between primary tumors with and without distant metastases did not demonstrate a statistically significant difference (p=0.32; not shown).

### Correlation of P2RY12 overexpression with overall survival

For survival analysis, Kaplan-Meier plots were created, and log-rank testing was applied (Figure 6). Samples were dichotomized into high and low P2RY12 expression according to the median of the OD values of the entire cohort (OD 0.141). Of the 478 biomarker-evaluable patients, 366 (76.6%) had valid follow-up information and were included (88 deaths within 5 years; median follow-up 3.3 years). Overall survival was analyzed for the whole cohort and, separately, for the subgroups LUAD and LUSC, and was visualized in Kaplan–Meier curves. For the entire cohort, patients with high P2RY12 protein expression showed a tendency towards better 5-year overall survival compared to those with low protein expression, although the difference was not statistically significant (p=0.31 by log-rank test; HR for low vs. high expression 1.24, 95% CI 0.82–1.89; n=366). Among patients with LUAD, those with high P2RY12 expression also showed a tendency towards better overall survival than those with low expression, without reaching statistical significance (p=0.19 by log-rank test; HR 1.51, 95% CI 0.81–2.81; n=225, 50 deaths). Similarly, in the LUSC group, overall survival was numerically higher in patients with high P2RY12 expression than in those with low expression, but the difference was not statistically significant (p=0.14 by log-rank test; HR 2.02, 95% CI 0.79–5.20; n=141, 38 deaths). Of note, only 15 LUSC patients fell into the low-expression group. Sensitivity analyses using histology-specific median cut-offs (LUAD p=0.26; LUSC p=0.16) or the visual score (moderate/high vs. absent/low; p=0.76) did not show significant differences either. Patients without follow-up information were more often LUAD (85.5% vs. 61.5%; p<0.001) and had slightly lower staining intensities (median OD 0.13 vs. 0.15; p=0.020), whereas age, sex, pT, pN, grading, and UICC stage did not differ.

**Figure 6.**
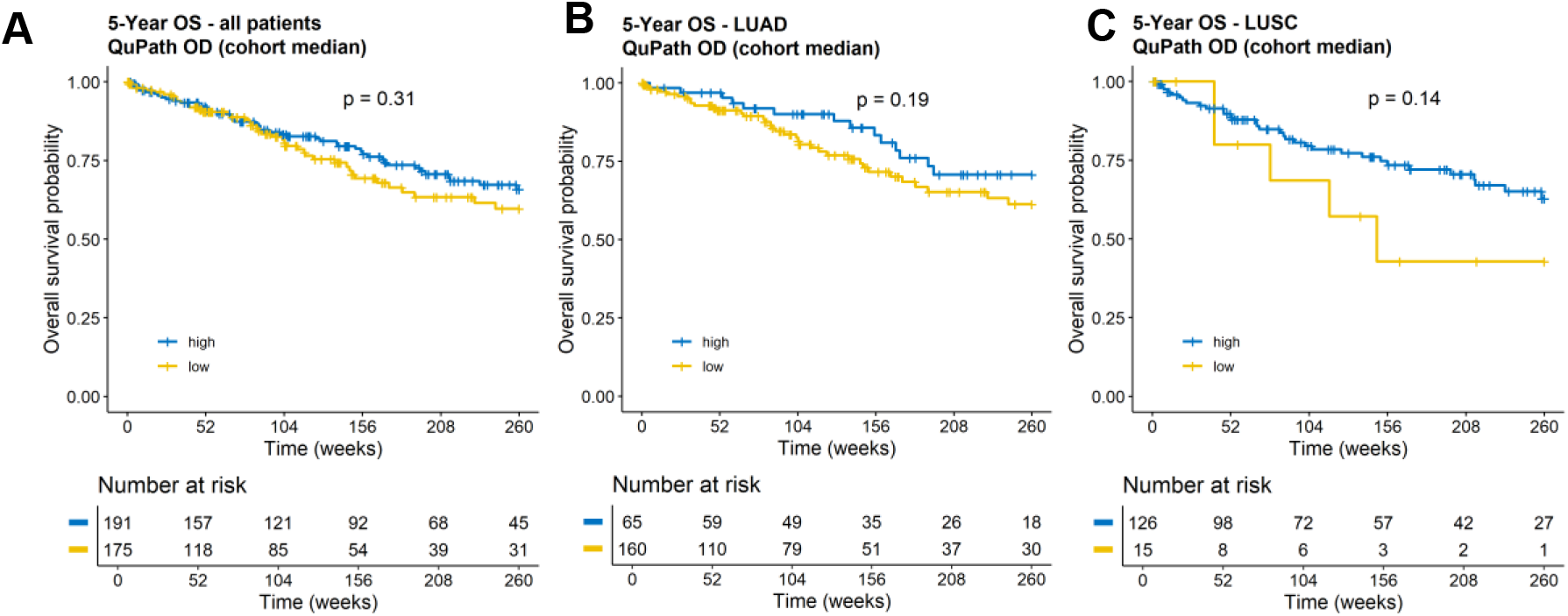
Kaplan-Meier graphs with p-values acquired from log-rank tests of (A) the entire cohort and separately for the subgroup (B) LUAD and (C) LUSC.

### Correlation of P2RY12 expression with further clinico-pathological characteristics

In the overall cohort of primary tumors (n=478), samples were stratified into high (n=239) and low (n=239) expression groups according to the median of the OD values. Staining results were correlated with TNM-Stage, UICC-Stage, grading, histologic subtype, age, and sex (Table 3; Fisher’s exact test for categorical and Wilcoxon rank-sum test for continuous variables). The distribution of T-Stage (T1–2 vs. T3–4) did not differ significantly between groups (p=0.371), with early-stage tumors predominating (69.3%). Nodal involvement was significantly more frequent in the low-expression group (36.3%) compared to the high-expression group (22.4%) (p=0.002). No significant differences were observed in M-Stage (p=0.221), UICC8-Stage (p=0.724), or UICC9-Stage (p=1.000). Tumor grading showed a balanced distribution between G1–2 and G3–4 (p=0.168). As already pointed out, histological subtype distribution differed markedly (p<0.001), with LUSC more prevalent in the high-expression group (59.5%) and LUAD predominating in the low-expression group (93.3%). Patients’ age was similar between groups (mean ± SD: high 67.9 ± 8.5 vs. low 66.4 ± 8.9 years, p=0.124), as was sex distribution (p=0.116).

**Table 3.** Correlation of P2RY12 expression with further clinico-pathological characteristics.

| Variable | P2RY12 Expression |  | Total (*n*=478) | *p* Value |
|---|---|---|---|---|
|  | high (*n*=239) | low (*n*=239) |  |  |
| **T-Stage** |  |  |  | 0.371 |
| - Missing | 1 | 1 | 0 |  |
| - T(1-2) | 160 (67.2%) | 170 (71.4%) | 330 (69.3%) |  |
| - T(3-4) | 78 (32.8%) | 68 (28.6%) | 146 (30.7%) |  |
| **N-Stage** |  |  |  | 0.002 |
| - Missing | 16 | 24 | 40 |  |
| - N(-) | 173 (77.6%) | 137 (63.7%) | 310 (70.8%) |  |
| - N(+) | 50 (22.4%) | 78 (36.3%) | 128 (29.2%) |  |
| **M-Stage** |  |  |  | 0.221 |
| - Missing | 172 | 205 | 377 |  |
| - M(-) | 64 (95.5%) | 30 (88.2%) | 94 (93.1%) |  |
| - M(+) | 3 (4.5%) | 4 (11.8%) | 7 (6.9%) |  |
| **UICC8 Stage** |  |  |  | 0.724 |
| - Missing | 11 | 13 | 24 |  |
| - UICC8(I-III) | 225 (98.7%) | 222 (98.2%) | 447 (98.5%) |  |
| - UICC8(IV) | 3 (1.3%) | 4 (1.8%) | 7 (1.5%) |  |
| **UICC9 Stage** |  |  |  | 1.000 |
| - Missing | 30 | 17 | 47 |  |
| - UICC9(I-III) | 206 (98.6%) | 219 (98.6%) | 425 (98.6%) |  |
| - UICC9(IV) | 3 (1.4%) | 3 (1.4%) | 6 (1.4%) |  |
| **Grading** |  |  |  | 0.168 |
| - Missing | 0 | 2 | 2 |  |
| - G(1-2) | 134 (56.1%) | 117 (49.4%) | 251 (52.7%) |  |
| - G(3-4) | 105 (43.9%) | 120 (50.6%) | 225 (47.3%) |  |
| **Histology** |  |  |  | < 0.001 |
| - Missing | 2 | 0 | 2 |  |
| - LUSC | 141 (59.5%) | 16 (6.7%) | 157 (33.0%) |  |
| - LUAD | 96 (40.5%) | 223 (93.3%) | 319 (67.0%) |  |
| **Age** |  |  |  | 0.124 |
| - Missing | 2 | 1 | 3 |  |
| - Mean (SD) | 67.867 (8.507) | 66.431 (8.911) | 67.146 (8.733) |  |
| - Range | 43.992 - 86.803 | 36.208 - 85.115 | 36.208 - 86.803 |  |
| **Sex** |  |  |  | 0.116 |
| - Female | 94 (39.3%) | 112 (46.9%) | 206 (43.1%) |  |
| - Male | 145 (60.7%) | 127 (53.1%) | 272 (56.9%) |  |

The multivariable Cox model included 330 patients and 76 deaths. After adjustment for age, sex, histology, nodal status, and tumour grade, with stratification by pT stage, low P2RY12 expression was not independently associated with overall survival (HR, 1.24; 95% CI, 0.69–2.24; p=0.47). No covariate was significantly associated with overall survival. In a model including pT stage as a covariate, the proportional hazards assumption was violated for pT stage (p=0.029; global test p=0.022), which is why the primary model was stratified by pT stage. In the stratified model, the global test did not provide statistically significant evidence against the proportional hazards assumption (p=0.058). Sensitivity analysis using multiple imputation yielded a similar estimate (HR, 1.49; 95% CI, 0.85–2.63; p=0.16). The model showed limited discrimination (concordance index, 0.62). With 76 events and six estimated parameters, the model had 12.7 events per parameter; variance inflation factors were below 2.3 for all covariates. When P2RY12 expression was entered as a continuous variable, the HR per 0.1 OD unit was 1.16 (95% CI, 0.91–1.49; p=0.24). Additional stratification by histology yielded a similar estimate (HR, 1.22; 95% CI, 0.68–2.20), and in histology-specific models the HR per 0.1 OD unit was 0.92 (95% CI, 0.43–1.96) in LUAD and 1.21 (95% CI, 0.93–1.57) in LUSC. The wide confidence intervals are compatible with both no effect and a moderate effect of P2RY12 expression on overall survival.

### P2RY12 Expression in the Differential Diagnosis of NSCLC Subtypes

Since P2RY12 expression was significantly higher in LUSC than in LUAD tissue (Table 2), we evaluated its discriminatory performance in distinguishing between these two NSCLC subtypes. Receiver operating characteristic (ROC) curve analysis demonstrated excellent discrimination, with an area under the curve (AUC) of 0.916 (95% CI 0.884–0.949 [DeLong]; 0.881–0.946 [stratified bootstrap]; n = 476; 157 LUSC, 319 LUAD) (Figure 7). The optimal cut-off, determined using the Youden index, was a QuPath optical density (OD) value of 0.196, yielding a sensitivity of 80.9% (95% CI 73.9–86.7%), a specificity of 94.0% (95% CI 90.9–96.4%), a positive predictive value of 87.0% (95% CI 80.4–92.0%), a negative predictive value of 90.9% (95% CI 87.3–93.8%), and an accuracy of 89.7% (95% CI 86.6–92.3%); the bootstrap 95% interval of the cut-off was 0.163–0.231. Internal validation showed little optimism: the bootstrap-corrected sensitivity and specificity were 79.7% and 93.5%, respectively, and repeated five-fold cross-validation yielded a mean sensitivity of 79.2% and a mean specificity of 94.2%. For visual scoring (moderate/high vs. absent/low), sensitivity was 78.3% (95% CI 71.1–84.5%) and specificity 98.4% (95% CI 96.4–99.5%). These findings indicate strong discriminatory performance within the investigated cohort, although independent validation is required.

**Figure 7.**
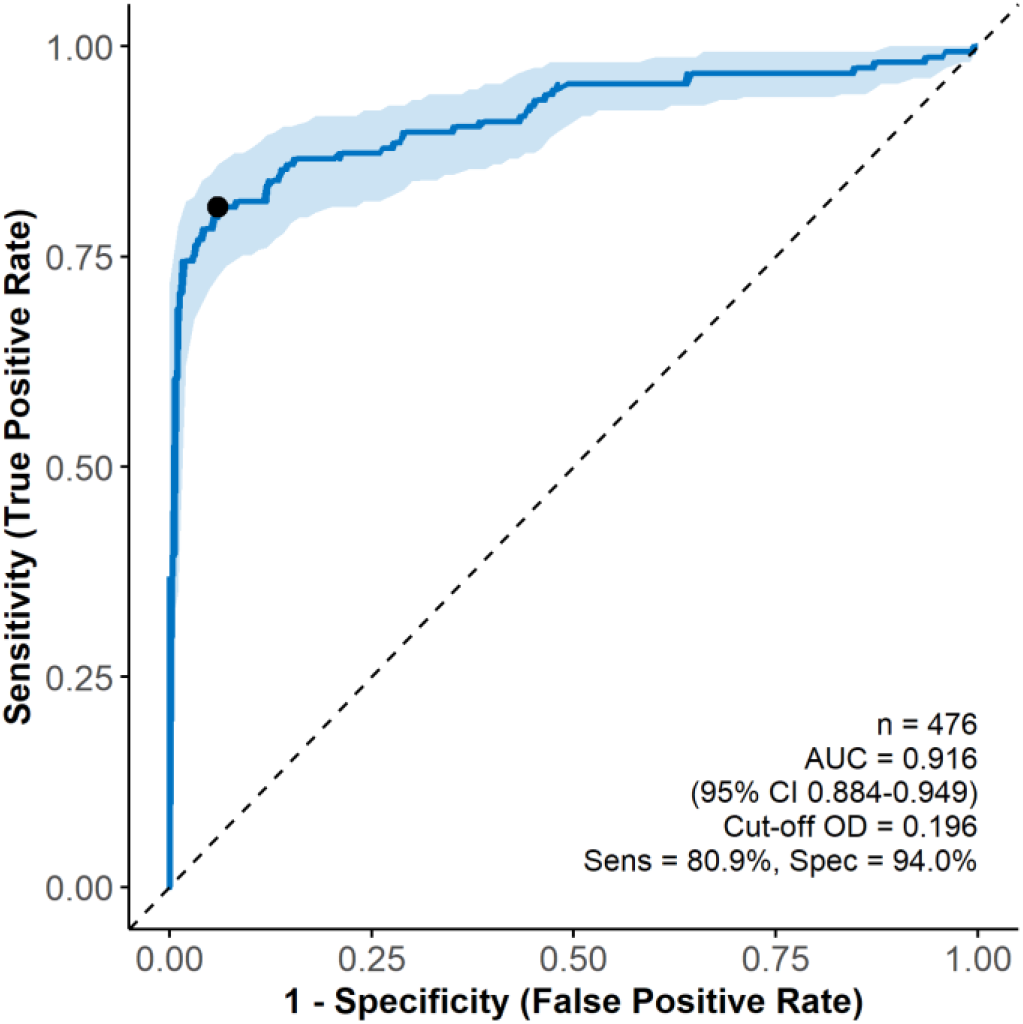
Receiver-operating-characteristic curve showing the discriminatory performance of P2RY12 staining intensity for differentiating LUAD from LUSC.

## Discussion

Despite promising advances in the treatment of NSCLC, there remains an urgent need to detect novel biomarkers with prognostic and therapeutic relevance. Among the two most common subtypes of NSCLC, patients with LUAD have benefited substantially from the advent of molecular targeted therapies, whereas progress in the treatment of LUSC has been limited (Howlader et al., 2020).

The purinergic receptor P2RY12 has been investigated in various malignancies, yet its role in NSCLC remains incompletely understood.

### P2RY12 expression pattern

To date, few studies have addressed P2RY12 protein expression in NSCLC. To our knowledge, the present study represents the first large-scale analysis of P2RY12 protein expression in a cohort of NSCLC, including both LUAD and LUSC, as well as metastatic lesions. We observed that P2RY12 expression was significantly higher in malignant than in benign lung tissue, except for benign bronchial epithelium, in which strong basal cell staining was detected, as exemplified in Figure 3i.

Whereas benign alveolar lung tissue displayed negligible P2RY12 expression (Figure 3h), malignant tissue exhibited variable staining intensities, ranging from absent to strong in LUSC and predominantly absent to weak in LUAD (Figure 3). These data suggest a potential role of P2RY12 in NSCLC pathogenesis, especially for LUSC.

The literature provides only partial agreement with these findings. Yu et al. reported an upregulation of P2RY12 mRNA in 352 LUAD samples derived from the Cancer Genome Atlas (TCGA) and performed immunohistochemistry on 37 LUAD specimens with corresponding adjacent benign tissues (Yu et al., 2021). They found lower protein expression in LUAD than in adjacent non-tumor samples. At first glance, this result might appear to contradict our own. However, the authors do not specify how they measured P2RY12 protein expression, nor do they explain in detail exactly which cells or which compartment of the lung the term ‘adjacent tissue’ refers to. We found negligible expression in pneumocytes but observed expression in respiratory epithelium (Figure 3h and i), which could in principle also be adjacent to cancer tissue. The authors refer solely to LUAD. In our subcohort of LUAD, we found, in line with this, predominantly (98%) low expression of P2RY12 (314 out of 319; Table 2). LUSC, however, for which we found substantial expression results, was not included in the cohort of the cited study. The functional experiments from Yu et al. suggested that P2RY12 overexpression suppressed proliferation and migration of LUAD cells, which is consistent with the numerically, but not significantly, higher survival probabilities in patients with high compared to low P2RY12 expression in our survival analyses (Figure 6).

Regarding P2RY12 expression in benign bronchial epithelial tissue, there is scarce information in the literature. Tsuya et al. found that P2Y receptor subtypes, including P2RY12, were expressed in immortalized human airway epithelial cell lines (16HBE cells) and identified P2RY12 as a mediator of ATP-induced enhancement of airway epithelial barrier integrity (Tsuya et al., 2026). Our observation of P2RY12 expression on benign bronchial epithelial tissue might suggest a possible role of P2RY12 in the earliest stages of LUSC development from stem cells/progenitor cells, ranging from squamous metaplasia and dysplasia to invasive squamous cell carcinoma. A review (Sánchez-Danés & Blanpain, 2018) reports on a study (Ferone et al., 2016) illustrating that basal and secretory cells of bronchial epithelium (as well as AT2 cells in alveoli) represent different possible cells of origin for LUSC in mice. Kim et al. investigated copy number variation (CNV) frequencies for pharmacogenes using the TCGA dataset and found that P2RY12 and P2RY1 had the highest frequency for gene gain in LUSC (Kim et al., 2015), which might point in that direction. This hypothesis would further support the possibility, discussed below, that P2RY12 might play a role as a diagnostic marker in LUSC, which warrants further investigation. However, our data are purely observational and thus not suitable for drawing conclusions that P2RY12 might play an etiological role in LUSC origin.

By comparing histological subtypes, we observed significantly higher P2RY12 expression in LUSC than in LUAD (p<0.001) (Table 2). To the best of our knowledge, no previous studies have directly compared P2RY12 protein expression between these two NSCLC subtypes. The inclusion of both histological subtypes in our comprehensive analysis makes our results novel.

### P2RY12 Expression in metastatic lesions

Since metastatic tissues are infrequently biopsied, large-scale analyses remain relatively scarce. In this regard, our study population provides valuable insights into dynamic changes in P2RY12 expression during tumor progression. Mean expression levels were significantly higher in LUSC compared with LUAD, lymph node metastases, and distant metastases. By contrast, no significant differences were observed between LUAD primaries and LNM or DM (Figure 5A), although markedly different sample sizes across the groups might influence these results (476 primary tumors, 34 lymph node metastases, and 42 distant metastases).

In paired specimens (n=38), metastatic deposits demonstrated a significant reduction in P2RY12 staining intensity relative to their primaries (p=0.025; Figure 5B). One reason could be that protein expression generally differs between primary tumors and metastases, probably in the context of tumor progression or temporal heterogeneity, as is known for programmed death ligand 1 (Wu et al., 2022). Another reason could lie in a context-dependent regulation of P2RY12 expression. Whereas platelets constitutively express high levels of P2RY12 as a prerequisite for aggregation, glioma and breast tumor cells have been shown to regulate P2RY12 expression dynamically in response to environmental stressors, such as serum deprivation or cytotoxic therapy (Czajkowski et al., 2004; Sarangi et al., 2013). Beyond such context-dependent changes in tumor cells, platelet/tumor cell signaling appears to involve distinct pathways, including down-regulation of SYK and inhibition of the PI3K–AKT signaling cascade (Song et al., 2025). However, both hypotheses are merely speculation, and our data are not sufficient to substantiate any closer connections in this regard.

Direct comparisons are of limited validity, and broad conclusions regarding a systematic down-regulation of P2RY12 in metastases cannot be drawn from our results. Most prior investigations have focused on the role of P2RY12 in platelets and tumor progression. Our findings underscore the need for further research to delineate the function of P2RY12 within tumor cells themselves.

### Concordance of evaluation methods

To ensure that our subjective visual evaluation and objective quantitative computer-based evaluation are in line with the literature (Jagomast et al., 2022), we compared subjective visual scoring with objective, computer-assisted quantitative evaluation. A strong correlation between the two methods was observed, supporting the validity of the classification in a low- and high-expressing group (Figure 4). For subsequent analyses, we therefore relied on the objective quantitative values.

DAB/AEC-based immunohistochemistry is widely used to visualize protein expression, with staining intensity serving as a surrogate for expression levels (Wen et al., 2024). However, the method is limited by dependence on technical variables, subjective scoring, and loss of information through categorical classification, rendering it semi-quantitative rather than fully quantitative (Ohkuma et al., 2023; Rizzardi et al., 2012). Automated image analysis seeks to overcome these constraints by providing more objective and reproducible measurements.

### Correlation of P2RY12 expression with survival

Kaplan–Meier analysis revealed no statistically significant association between P2RY12 protein expression and 5-year overall survival neither in the entire NSCLC cohort (log-rank p = 0.31) nor in the LUAD (p = 0.19) nor LUSC (p = 0.14) subgroups (Figure 6). Nevertheless, patients with high P2RY12 expression showed numerically higher survival probabilities across all three analyses. Multivariable Cox regression likewise did not identify P2RY12 expression as an independent prognostic factor. These findings do not establish a prognostic role for P2RY12 in NSCLC, although a potential association cannot be excluded.

Our findings may also be considered in the context of previous studies investigating P2RY12 expression in lung cancer. Yu et al. reported an association between reduced P2RY12 protein expression and poorer overall survival in a small LUAD cohort (n = 37) (Yu et al., 2021). Although the direction of the numerical survival differences in our study was similar, we could not confirm a statistically significant association in our larger LUAD cohort. Differences in cohort characteristics, biomarker assessment, and expression thresholds may contribute to these findings. Furthermore, our study extends the investigation of P2RY12 expression to LUSC, for which prognostic data remain limited.

Interestingly, large-scale epidemiological studies have demonstrated that LUAD generally has a more favorable prognosis than LUSC. In a retrospective analysis of 48,296 NSCLC patients, Wang et al. reported a better stage-specific 5-year overall survival in all tumor stages for LUAD compared with LUSC (B.-Y. Wang et al., 2020). In our cohort, however, P2RY12 expression was substantially higher in LUSC than in LUAD, while patients with high P2RY12 expression exhibited numerically higher survival probabilities. Notably, this pattern was also observed within the histologically stratified LUAD and LUSC subgroups, suggesting that the observed differences may not be solely attributable to histological composition. Although these observations are consistent with a possible favorable association between P2RY12 expression and survival, they did not reach statistical significance and cannot establish an independent prognostic effect.

The biological role of P2RY12 in lung cancer may depend on its cellular context. Previous preclinical studies have implicated platelet P2RY12 signaling in tumor progression and metastasis (Gresele et al., 2018; Mege et al., 2019). These observations do not necessarily reflect the consequences of P2RY12 expression within tumor cells. Increased extracellular ADP in the tumor microenvironment (Di Virgilio et al., 2018) may promote platelet activation and influence purinergic signaling, potentially resulting in distinct effects across cellular compartments. Whether tumor-cell P2RY12 expression reflects altered signaling activity or other characteristics of the tumor microenvironment remains uncertain. Further functional studies are needed to clarify these mechanisms and determine whether P2RY12 has prognostic relevance in NSCLC.

### Clinico-pathological correlations of P2RY12 expression

No significant associations were observed between P2RY12 expression and patient sex, age, tumor grade, T-stage, M-stage, or UICC-stage. In contrast, a significantly higher frequency of nodal involvement was detected in the low-expression group (p=0.002; Table 3). This observation lends some plausibility to our findings, as it is in line with the observed survival trend demonstrating a trend toward poorer overall survival for the P2RY12 low expressing group. However, given that we did not identify any statistically significant associations with other clinicopathological parameters, we cannot exclude the possibility that this finding represents a chance observation or an epiphenomenon. Therefore, we believe that this result should be interpreted with caution and not be overestimated.

This observation appears further inconsistent with preclinical murine and *in vitro* models, in which genetic deficiency or pharmacologic inhibition of platelet P2RY12 reduced metastatic spread (Y. Wang et al., 2013). Conversely, high P2RY12 expression has been shown to suppress migration and proliferation of LUAD cells (Yu et al., 2021). The results we have obtained from FFPE material are naturally not directly comparable with *in vitro* analyses, of which there are few in the literature.

### Diagnostic Potential of P2RY12 expression

P2RY12 demonstrated strong discriminatory capacity between LUAD and LUSC, with an area under the receiver-operating-characteristic curve (AUC) of 0.916. This diagnostic accuracy is comparable to established immunohistochemical markers such as TTF-1, Napsin A, p63, and p40 (Grover et al., 2024), suggesting potential utility as a diagnostic adjunct.

The precise histologic and subsequent molecular classification is essential for treatment stratification in NSCLC, and P2RY12 may thus complement existing diagnostic markers. This would also be significant for metastatic tissue. In particular, P2RY12 might improve the accuracy of NSCLC subclassification in cases with equivocal expression of conventional markers. This is of considerable clinical relevance, as accurate diagnosis is a prerequisite for selecting the most effective personalized therapy in order to substantially influence outcomes in patients with NSCLC (M. Wang et al., 2021). As this study is the first to investigate P2RY12 expression in a cohort comprising both adenocarcinomas and squamous cell carcinomas, it would be important to test this hypothesis further in larger validation cohorts of NSCLC. Moreover, other organs that may harbour both adenocarcinomas and squamous cell carcinomas (e.g. cervix uteri, oesophagogastric junction) should be analyzed for the diagnostic value of P2RY12. Due to several immunohistochemical markers of squamous differentiation, such as p40, p63, and CK5/14, are already available, it is admittedly questionable whether an additional marker of squamous differentiation is essentially necessary in routine diagnostic. Nevertheless, our observation regarding the potential diagnostic utility of P2RY12 is of considerable interest.

### Limitations

The distribution of T-stages within our cohort was skewed toward early-stage disease, with a dominant part of tumors classified as T1 or T2 (68.9%). This does not reflect the general stage distribution reported in population-based studies (Goldstraw et al., 2016) and is explained by our reliance on surgically resected tissue for TMA generation. Patients undergoing primary surgery typically present with lower T-stages, while those with advanced stages may not be surgical candidates or may undergo surgery only after neoadjuvant therapy (Hendriks et al., 2024). Importantly, no specimens obtained after neoadjuvant treatment were included in our study.

Another limitation is the lack of validation in an independent cohort. Previous studies have primarily investigated P2RY12 at the mRNA level or involved significantly smaller or compositionally different cohorts. No comparative data on P2RY12 protein expression in NSCLC are currently available in the literature, underscoring the novelty of our findings but also highlighting the need for independent replication.

Another important methodological consideration is the dichotomization of continuous biomarker expression values. As highlighted by Polley and Dignam, categorization using median-based cut-offs may result in substantial information loss and reduce the ability to detect associations with clinical outcomes (Polley & Dignam, 2021). Thus, the absence of statistically significant differences in Kaplan–Meier analyses does not necessarily exclude a potential prognostic relationship. Nevertheless, such findings require cautious interpretation, particularly in the context of exploratory subgroup analyses, limited numbers of events, and potential confounding by established prognostic factors. Analyses incorporating continuous expression measurements and validation in independent cohorts are therefore warranted.

Even though our results demonstrate excellent discriminatory performance of P2RY12 expression for distinguishing LUAD from LUSC (AUC 0.916), this finding requires external validation in larger, prospectively collected cohorts. Specifically, diagnostically challenging cases, such as poorly differentiated tumors, adenosquamous carcinomas, and small biopsy or cytology/cell block specimens, should be included to assess whether P2RY12 can serve as a reliable immunohistochemical marker for squamous differentiation.

There are *in vitro* data in the literature showing that the P2RY12 antagonist Ticagrelor significantly inhibits the proliferation, migration, and invasion of several cell lines derived from lung adenocarcinomas (Song et al., 2025). A significant limitation, therefore, is that our study does not include any functional analyses. Especially for LUSC cell lines, this would be of particular interest, based on our findings. This should therefore be investigated in further studies.

Another limitation lies in the lack of detailed patient histories, which include information regarding cardiovascular diseases and therapy with P2Y12 inhibitors to prevent thrombocyte aggregation. Consequently, no correlation between an appropriate treatment and the observed expression patterns can be established here. This question warrants systematic evaluation in future studies with comprehensive clinical data, although results of *in vitro* analyses using cancer cell lines cannot, of course, be directly extrapolated to processes in the human body. A further limitation of our study is the absence of progression-free survival data, which precluded assessment of P2RY12 as a prognostic biomarker in this context.

*<u>In summary</u>*, the expression patterns and clinical relevance of P2RY12 in NSCLC remain insufficiently characterized. Previous studies support a role of P2RY12 in tumor progression and metastasis, and experimental data across cancer entities suggest potential therapeutic relevance (Song et al., 2025). Our analyses in a large NSCLC cohort demonstrated differential P2RY12 protein expression between LUAD and LUSC, and numerically, but not significantly, lower survival probabilities in patients with reduced expression.

Given its reliable immunohistochemical detectability, simple classification, and strong discriminatory capacity between NSCLC subtypes, P2RY12 might serve as a new diagnostic biomarker for LUSC. Incorporation into routine diagnostics appears feasible. Given the potential diagnostic utility for squamous differentiation of this biomarker demonstrated in our study, it would be essential to further investigate its prognostic relevance in larger cohorts, especially of LUSC patients, particularly given the relatively limited number of squamous cell carcinoma cases included in our study.

P2RY12 is known as an actionable target, and P2RY12 antagonists are in clinical use for cardiovascular diseases. That is why, despite the lack of independent prognostic significance, P2RY12 remains a promising candidate for further investigation as a potential therapeutic target in NSCLC, especially considering the availability of clinically approved P2RY12 antagonists that may offer opportunities for drug repurposing. Beyond further evaluating the prognostic relevance of P2RY12 in larger NSCLC/LUSC cohorts, future studies should investigate its potential as a therapeutic target and explore whether P2RY12 expression may serve as a predictive biomarker for response to targeted therapies.

## Acknowledgments

We thank Christian Rosero for the excellent technical support.

## Data availability statement

The raw data supporting the conclusions of this article will be made available by the authors, without undue reservation.

## Ethics statement

The study was conducted in accordance with the Declaration of Helsinki, and the protocol was approved by the local ethics council at the University of Lübeck (file number 16-277 and 16-278).

## Author contributions

TG and CK planned the research project. TJ performed and visualized the statistical analysis. HR, LE, CH, F-OP, TT, SB, MS, MR, TO, SW provided patients’ follow-up data. CK, HR, and TG wrote the manuscript, and TJ, LE, CH, F-OP, TT, JK, DD, SB, MS, MR, TO, SW, VM, and DN revised it. All authors have read and agreed to the published version of the manuscript.

## Conflict of interest

CK has given paid lectures for AstraZeneca, MSD, Histoserve/Diapath and Boehringer Ingelheim.

The (remaining) authors declare that the research was conducted in the absence of any commercial or financial relationships that could be construed as a potential conflict of interest.

The above-mentioned companies had no influence on the study design, acquisition of data, or writing of the manuscript.

## Abbreviations

AEC: 3-amino-9-ethylcarbazole;
ATP: adenosine triphosphate;
AUC: area under the curve;
CNV: copy number variation;
DAB: 3,3′-diaminobenzidine;
DM: distant metastasis;
EMT: epithelial-mesenchymal transition;
FFPE: Formalin-Fixed and Paraffin-Embedded;
HRP: Horseradish Peroxidase;
IHC: immunohistochemistry;
LLC: Lewis Lung Carcinoma;
LNM: lymph node metastasis;
LUAD: pulmonary adenocarcinoma;
LUSC: pulmonary squamous cell carcinoma;
mRNA: messenger ribonucleic acid;
NK cells: natural killer cells;
NOS: not other specified;
NSCLC: non-small cell lung cancer;
OD: optical density;
OS: overall survival;
P2RY12: platelet P2RY12 receptor;
QuPath: Quantitative Pathology;
ROC: receiver-operating-characteristic;
ROI: region of interest;
SCLC: small cell lung cancer;
TCGA: The Cancer Genome Atlas;
TMA: tissue microarray;
TNF-α: tumor necrosis factor alpha;
UICC: Union for International Cancer Control;
VEGF: vascular endothelial growth factor;
WHO: World Health Organization

## Supplement

**Supplement Figure 1.**
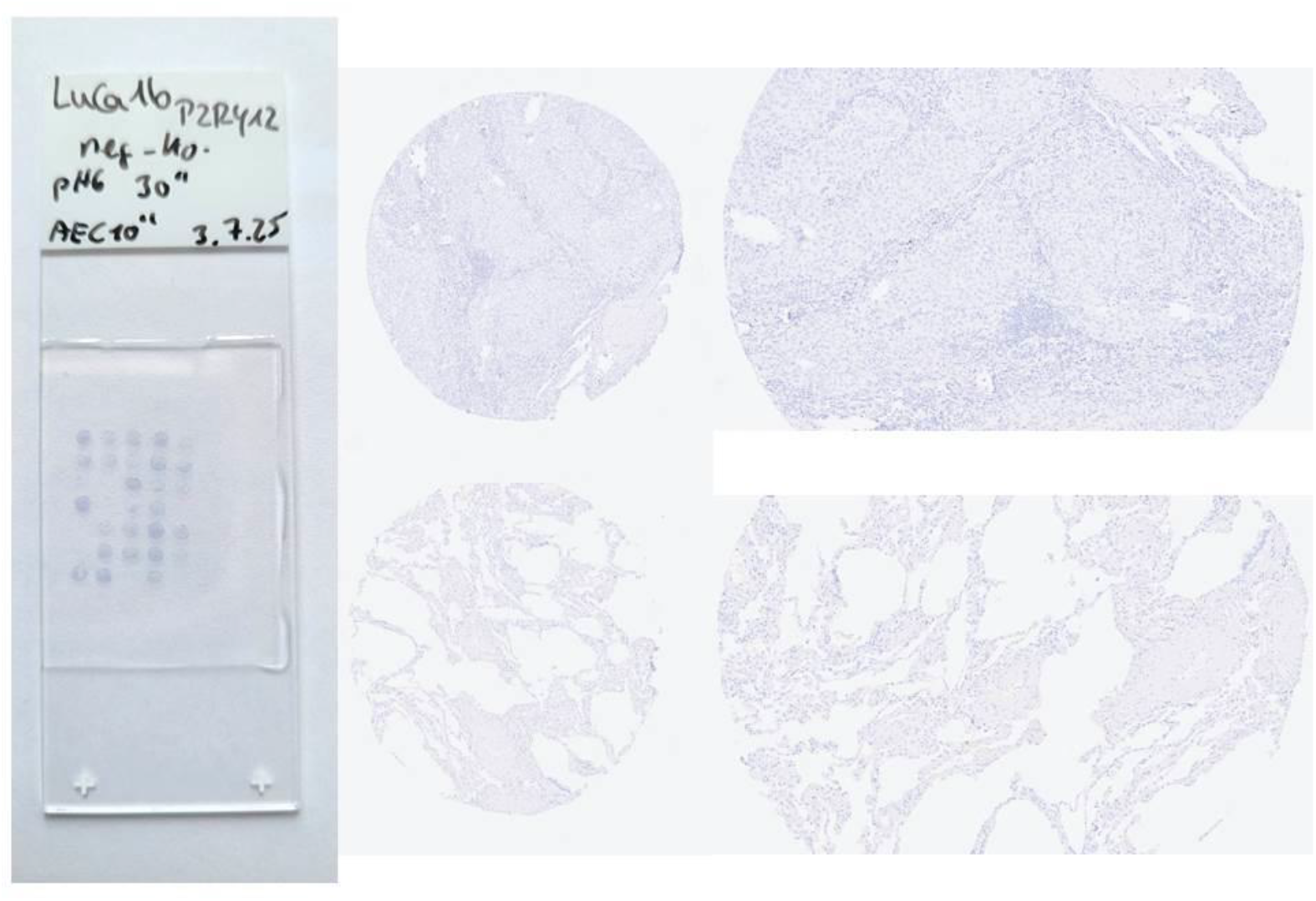
Pictures of the negative control without the primary P2RY12 antibody

